# SiCoDEA: a simple, fast and complete app for analyzing the effect of individual drugs and their combinations

**DOI:** 10.1101/2022.04.19.488737

**Authors:** Giulio Spinozzi, Valentina Tini, Alessio Ferrari, Ilaria Gionfriddo, Roberta Ranieri, Francesca Milano, Sara Pierangeli, Serena Donnini, Serenella Silvestri, Brunangelo Falini, Maria Paola Martelli

## Abstract

The administration of combinations of drugs is a method widely used in the treatment of different pathologies as it can lead to an increase in the therapeutic effect and a reduction in the dose compared to the administration of the single drugs. For these reasons, it is of interest to study combinations of drugs and in particular to determine whether a specific combination has a synergistic, antagonistic or additive effect, i.e greater, less than or equal to the effect expected by the sum of the individual drugs. For this purpose, various mathematical models have been developed, which use different methods to evaluate the synergy of a combination of drugs. Most of these methods are based on the Loewe Additivity Principle (Loewe et al., 1953), which has its key step in choosing the model used for predicting the effect of individual drugs. Creating a model for this purpose and calculating its parameters, however, requires a certain level of mathematical and programming knowledge or the use of commercial software. For this purpose, therefore, we have developed an open access and easy to use app that allows to explore different models and to choose the most fitting for the specific experimental data: SiCoDEA (Single and Combined Drug Effect Analysis, https://sicodea.shinyapps.io/shiny/). The data used to test SiCoDEA comes from cell line samples treated with different drug combinations and analyzed through metabolic or viability assays. SiCoDEA is developed through a Shiny interactive and easy-to-use interface (R based). There are five models taken into consideration for the analysis of single drugs and the calculation of combination index. The first is one of the most used, that of the median-effect; while the others are different forms of the log-logistic equation, with two, three and four parameters. The purpose of SiCoDEA is, on the one hand, to provide an easy-to-use tool for analyzing drug combination data and, on the other hand, also to have a view of the various steps and to offer different results based on the model chosen. An important prerequisite in analyzing drug combinations is in fact the dose-response curve calculated for individual drugs. For this purpose, SiCoDEA allows you to view the plots of the individual drugs both to evaluate the distribution of the calculated points and therefore identify any outliers, and to view the curve of the different models taken into consideration and evaluate which one best fits the data. A table showing all the R2 values for the five different models is created with the curve plot. In addition to the type of model, it is also possible to choose between two different normalization methods, one based on the maximum or minimum value and the other on the value calculated at drug concentrations equal to zero. For the chosen options, a plot is then created that shows the trend of the combination index for the different drug combinations and consequently whether it is synergy, antagonism or additivity. Finally, it is possible to export the results in single png files or in a summary report in pdf. SiCoDEA is an open-source app among the most complete and offers more functions even than the famous CompuSyn (Chou et al., 2010), as it allows you to analyze drug curves with different models, rather than just one, and it also allows the analysis of single drug curves.

## Motivation

Since 1965 when Emil Frei designed the first combinatorial regimen for acute leukemia [1] it became clear that remissions obtained with therapies based on single drugs were only temporary, and that the clinical responses achieved would be more durable when more agents were combined.

Over time, pursuing this approach, cancers that had previously been fatal such as acute lymphocytic leukemia, diffuse large B-cell lymphoma, Hodgkin’s lymphoma and testicular cancer became largely curable [2].

These days effective chemotherapies mostly involve combinations of two or more drugs allowing the targeting of tumor heterogeneity, feedback loops, dependencies, synthetic lethality, and the selective rise of therapy-resistant tumor clones.

In the last two decades, a new concept of cancer therapy, the targeted therapy, has emerged giving rise to several classes of cancer drugs that are designed to precisely block specific pathways that are relatively selective to the cells of distinct cancer types, inhibiting their growth or promoting their differentiation or death sparing healthy tissues. These new targeted therapies that include kinases inhibitors, receptor inhibitors, immunotherapies give promising results in combination with standard chemotherapeutics regimens [3, 4] and equally growing is the depth of molecular characterization in cancer.

With countless possibilities of combinations, dosing, scheduling, and repurposing, and finer targeting due to the deeper disease characterization the number of possible clinical trials vastly exceeds the number of patients [5].

If we take into account that just a 7% of anticancer drugs undergoing clinical trials proves to be effective [6], while oncology trials are, together with the cardiovascular, the most expensive [7]. It becomes even more clear that technological advances are urgently needed to make the design of combination treatments, necessary to improve clinical outcome for most patients, truly fruitful.

For these reasons, it is of interest to study combinations of drugs and to determine whether a specific combination has a synergistic, antagonistic, or additive effect, i.e., greater, less than or equal to the effect expected by the sum of the individual drugs.

The fundamental step is the definition of a Combination Index (CI) that allows to evaluate the effect of the two drugs used separately with respect to the combination. CI represents a value that indicates the distance of the observed response from the expected response and in particular it will indicate synergy if *CI* < 1, antagonism if *CI* > 1, and additivity if *CI* = 1. For this purpose, an *Effect-Based Strategy* or a *Dose-Effect-Based Strategy* can be used [8], two different approaches that compare the observed effect of the combination with the expected effect under the assumption of non-interaction predicted by a reference model. In the first case, a direct comparison is made between the effect of the individual drugs and the effect of the drugs on the combination, in the second case the calculation of the dose-effect curves for the individual drugs is used to calculate their expected values. In this second case, therefore, a further important step is the choice of the model to calculate the dose-effect curve. The Effect-Based Strategy include the *Response Additivity* model, the *Highest Single Agent* (HSA) model and the *Bliss Independence* model, while the Dose-Effect-Based Strategy include the *Loewe Additivity* model and the *Zero Interaction Potency* (ZIP) model.

### Response additivity model

Also known as the *Linear Interaction Effect*, the *Response Additivity* model [9] consists in comparing the effect of the combination and the effect obtained by adding that of the individual drugs at the same dose. It is therefore possible to obtain a CI from the ratio: 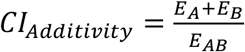 where *E_A_* is the effect of drug A alone, *E_B_* is the effect of drug B alone and *E_AB_* is the effect of drugs A and B in combination.

### Highest Single Agent model (HSA)

The *Highest Single Agent* model (HSA) or *Gaddum’s noninteraction* model [10] assumes that the expected effect of the combination is equal to the highest effect of the individual drug at the same dose as it has in the combination. Thus, a synergistic combination should produce an additional beneficial effect compared to what individual drugs alone can achieve. The CI is given by the difference between the effect of the combination at a given dose and the highest effect of one of the single drugs at that same dose: 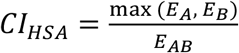.

### Bliss independence model

The *Bliss Independence* model [11] assumes a stochastic process in which two drugs produce their effect independently. Therefore, the expected effect of the combination can be calculated as the probability of two independent events: *E_A_* + *E_B_* – *E_A_ E_B_* where 0 ≤ *E_A_* ≤ 1 e 0≤ *E_B_* ≤ 1. The CI will be: 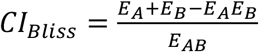.

### Loewe additivity model

The principle on which the *Loewe Additivity* model [12–14] is based is that to calculate the CI it is necessary to compare the doses of the drugs in combination with the doses of the individual drugs necessary to achieve the same effect. In this way, if the dose required for a single drug is lower than that in combination, we will have an antagonistic effect between the two drugs, if instead it is higher the effect will be synergistic.

The CI will be calculated as: 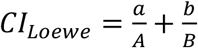 where *a* is the dose of drug A in the combination, *A* is the equivalent dose, i.e. the dose of drug A needed to achieve the same effect of the combination, *b* is the dose of drug B in the combination and *B* is the equivalent dose, i.e. the dose of drug B needed to achieve the same effect of the combination.

### Zero Interaction Potency (ZIP) model

The *Zero Interaction Potency* (ZIP) model [15, 16] combines the Loewe model and the Bliss model together assuming that in combination the two dose-effect curves do not change. It then uses the same calculation of two independent events of the Bliss model, but using the values calculated through the dose-effect curve as in the Loewe model. The CI will be: 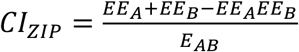 where *EE_A_* is the expected effect of drug A and *EE_B_* is the expected effect of drug B calculated based on the dose-effect curve.

#### Dose-response curve models

In the case of Dose-Effect-Based approaches, the choice of the model for the calculation of the doseresponse curve is equally important. Although, in fact, there are models that are more used than others, it is important to evaluate case by case, on the basis of the data under examination, which model is best suited.

The most commonly used model is the *Chou-Talalay Method* [17] which is based on the median-effect equation, derived from the mass-action law principle:

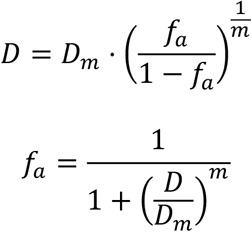

where *D* is the dose of interest, *D_m_* is the median-effect dose, *f_a_* is the fraction affected and *m* is the slope.

Another widely used model is the log-logistic one, which can use two, three or four parameters: with two parameters the minimum is set equal to zero and the maximum equal to one; with three parameters only one of the two is kept fixed (either the maximum or the minimum); finally, with four parameters there is no fixed value, but all four are calculated within the model (maximum, minimum, median-effect dose, *D_m_*, and slope, *m*). The following are the formulas for the four models:

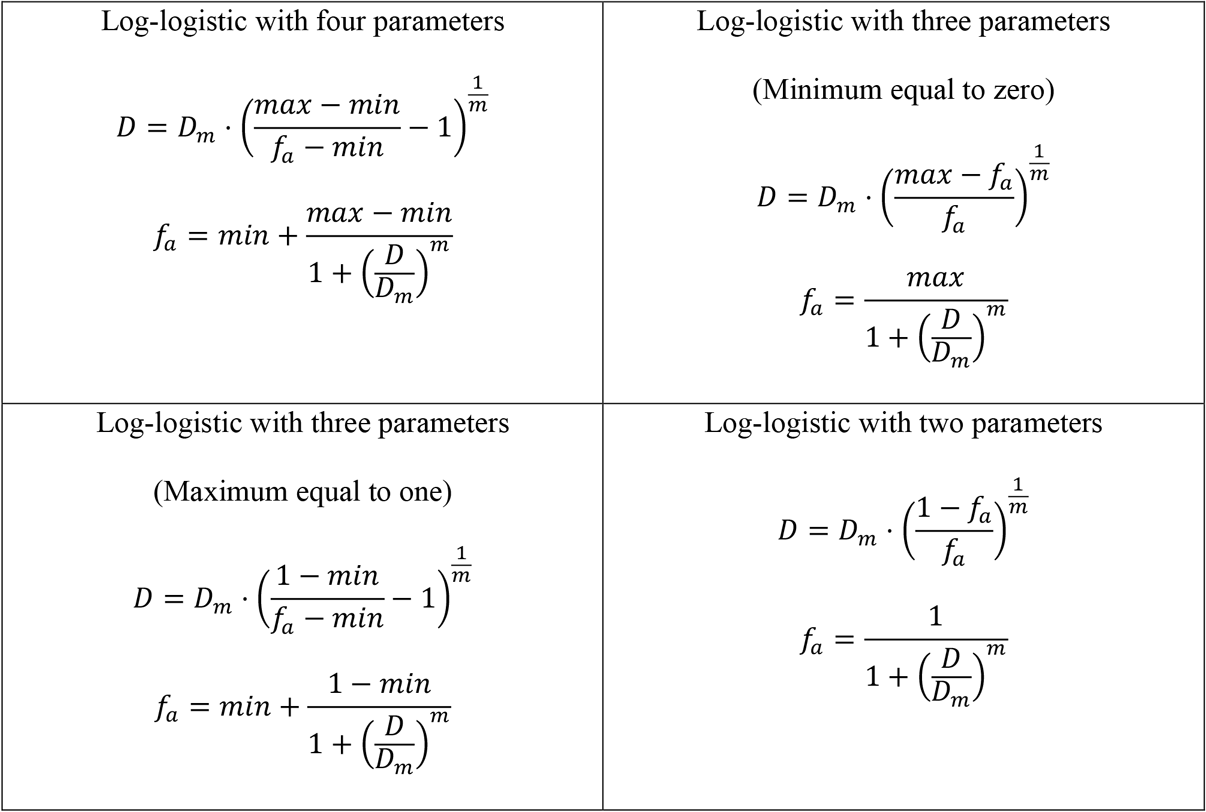

Each of these methods for calculating the CI has advantages and disadvantages based on the situation and the data being analyzed and for this reason it is of interest to provide a choice. Creating a model for this purpose and calculating its parameters, however, requires a certain level of mathematical and programming knowledge or the use of commercial software.

There are already several tools that analyze the interaction between drugs and the most famous or complete are CompuSyn [18], SynergyFinder Plus [19] and DDCV [20]. However, we have observed that, for an analysis that is as precise and flexible as possible, something is missing from each of these tools (Table 1). CompuSyn is a paid software that only works on Windows platforms, it also doesn’t offer many options to choose from and only allows analysis using the median-effect model. DDCV is another shiny app that can be found on the web that allows the analysis of drug combinations. It uses only the median-effect model for the calculation of the CI, without the possibility of choosing otherwise. Another fairly complete app that allows the analysis of drug combinations is SynergyFinder Plus, but, although it allows you to choose between different models for the calculation of the CI, it does not allow you to choose between different models of dose-response curve in the web version. In addition, none of these tools offer an analysis of the outliers with variable filter threshold, in order to refine the model.

**Table 1.** Options available in the various tools for drug combination analysis.

|  | CI models | Drug-response models | Open source | Report | Single drug analysis | Customable outlier analysis | Platform |
| --- | --- | --- | --- | --- | --- | --- | --- |
| **SynergyFinder Plus** | 4 | 1 | ✓ | ✓ | ✓ |  | Win/Mac/Linux |
| **CompuSyn** | 1 | 1 |  |  | ✓ |  | Win |
| **DDCV** | 1 | 1 | ✓ | ✓ |  |  | Win/Mac/Linux |

For this purpose, therefore, we have developed an open access and easy to use app that allows to explore different models and to choose the most fitting for the specific experimental data: SiCoDEA (Single and Combined Drug Effect Analysis).

## Methods

The validation of our novel SiCoDEA open-source app was performed on multiple AML cell lines: OCI-AML3, IMS-M2, OCI-AML2, SKM-1, MV-4-11, MOLM-13 cultured as recommended by cell line providers and previously described [21]. For drug data analysis, the AML cells were exposed to drugs or controls after single and combination drug treatment using compounds already in use in our laboratory.

The first step was the optimization of drug responsive curves for the drugs used as single agents.

Cells were seeded in 384-well plate at a concentration of 4 × 105 cells/ml in a volume of 22.5 μl/well and treated in triplicate with vehicle or 7 log-scale concentrations of each drug using the automized D300e Digital Dispenser (Tecan). 100 μM Etoposide treatment was used as positive control. After 48 hours, cell proliferation was assessed using the cell metabolism independent Cyquant Direct Cell Proliferation Assay (Life Technologies). Cells were incubated for 4 hours of with CyQuant reagent and then fluorescence signal of treated and non-treated control were measured in a Spark Microplate Reader (Tecan).

Data were normalized assuming 100% cell proliferation to untreated control and 0% proliferation to 100 μM Etoposide treatment.

Once drug responsive curves had been optimized, we proceeded to drug combinatorial treatment.

Again, cells were seeded in 384-well plates and treated in triplicate with an 8×8 matrix of log-scale drug concentrations ranging from 0 to the maximum effective dose for each drug. The IC50 of each drug was in the middle of the 7 drug doses. 48 hours later Cyquant reagent was added, and data were analysed in order to assess the CI.

SiCoDEA (available at https://sicodea.shinyapps.io/shiny/) is developed through a *Shiny* interactive and easy-to-use interface (R based); it provides both the simple calculation of the IC50 for different drugs and the calculation of the CI with the display of the respective plots. The utilized R packages are “shiny” [22], “shinyjs”[23], “plyr”[24], “car”[25], “drc”[26], “ggplot2”[27], “tidyr” [28], “gplots” [29], “outliers” [30], “scales” [31], “rlist” [32], “dplyr” [33]. A more detailed description of its use is available in *Additional File 1*.

## Results and discussion

The purpose of SiCoDEA is, on the one hand, to provide an easy-to-use tool for analyzing drug combination data and, on the other hand, also to have a view of the various steps and to offer different results based on the model chosen. An important prerequisite in analyzing drug combinations is in fact the dose-response curve calculated for individual drugs. Many of the existing tools, from the famous Compusyn to the most recent SynergyFinder Plus, involve the use of a single model in the calculation of the dose-response curve, but we have observed that there is not universally better model than the others: different data require different models. In cases such as that in Figure 1A, for example, the four-parameter log-logistic model appears to be the one with the greatest R^2^. In Figure 1B, however, we have instead a case in which the trend of the drug is better described by a three-parameter log-logistic model. Furthermore, the situation becomes complicated when two different drugs must be considered, as in the case of combinations. For this reason, SiCoDEA allows you to view the plots of the individual drugs with the curve of the different models taken into consideration and evaluate which one best fits the data. A table showing all the R^2^ values for the five different models is created with the curve plot. In addition to the type of model, it is also possible to choose between two different normalization methods, one based on the maximum or minimum value and the other on the value calculated at drug concentrations equal to zero.

**Figure 1.**
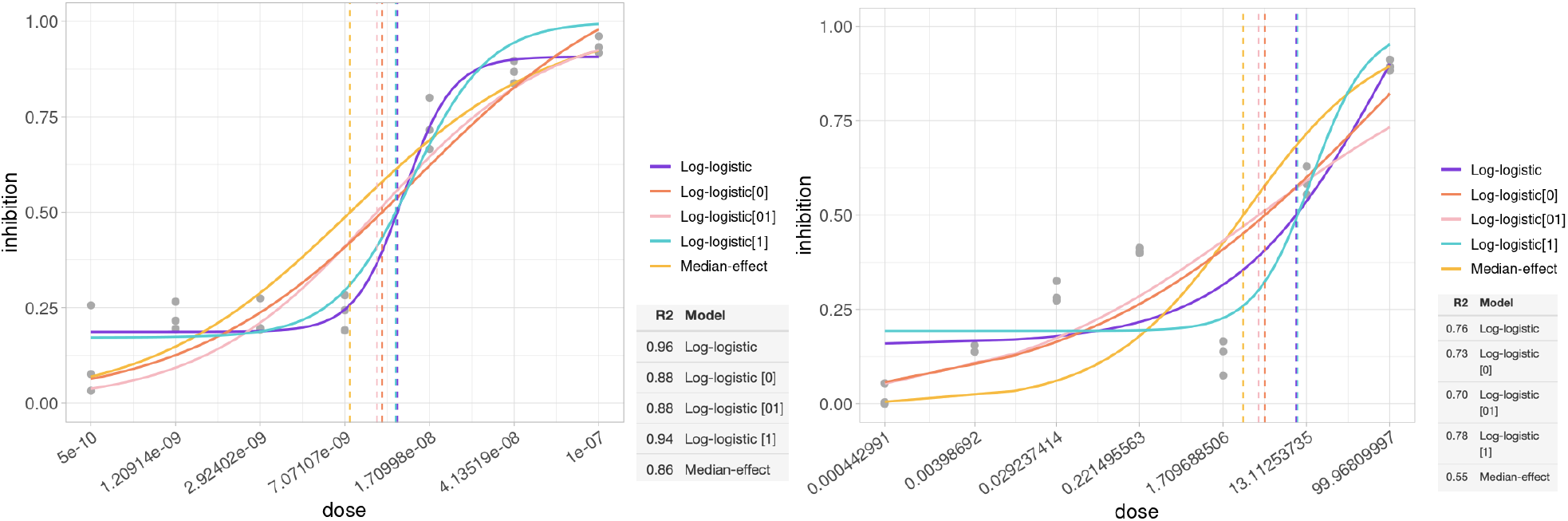
Dose-response curves applying five different models to the same data, the drug concentration is shown in the x axis and the proportion of cells that have undergone inhibition in the y axis. The lower right table shows the R^2^ value for each. In the first case (A) the model with the highest R^2^ and therefore best suited to the data is the log-logistics with 4 parameters, while in the second case (B) it is the log-logistics with 3 parameters with a maximum value set at 1.

Another element that can compromise the calculation of the model parameters, and therefore the result, is the presence of outliers (Figure 2A before outlier removal and figure 2B after outlier removal). SiCoDEA therefore allows you to choose the p-value threshold to use for the outlier calculation. In particular, a Grubbs’ test [34] is carried out on the basis of which a p-value equal to 0 means that no outlier, while a value equal to 1 means that a replicate will be eliminated for each dose concentration. By changing the threshold, you can immediately observe the consequences on the model to choose the right value to obtain the most reliable results.

**Figure 2.**
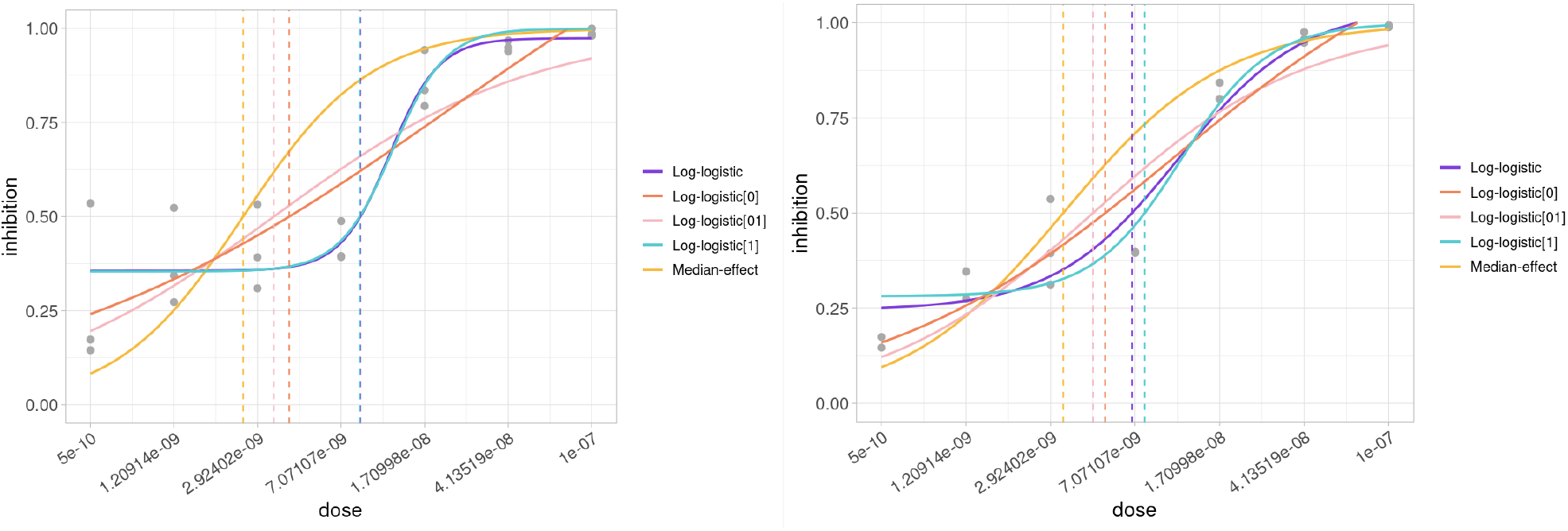
Dose-response curves showing models and data before (A) and after (B) outlier removal. The concentration of the drug is shown on the x axis and the proportion of inhibition is shown on the y axis.

For the chosen options, a plot is then created that shows the trend of the CI for the different drug combinations and consequently whether it is synergy, antagonism, or additivity. Finally, it is possible to export the results in single png files or in a summary report in pdf.

The app is divided into three tabs, each dedicated to a different analysis.

The first tab (Figure 3) is dedicated to single drug analysis and has the advantage of being able to analyze many drugs in a single file, it is sufficient to enter one drug per line, both in the dose file and in the response file. The plot shows the curves for all five models, as well as the line corresponding to the IC50 for each model. Based on the curves and the calculated R^2^, the model that best fits the data can be chosen.

**Figure 3.**
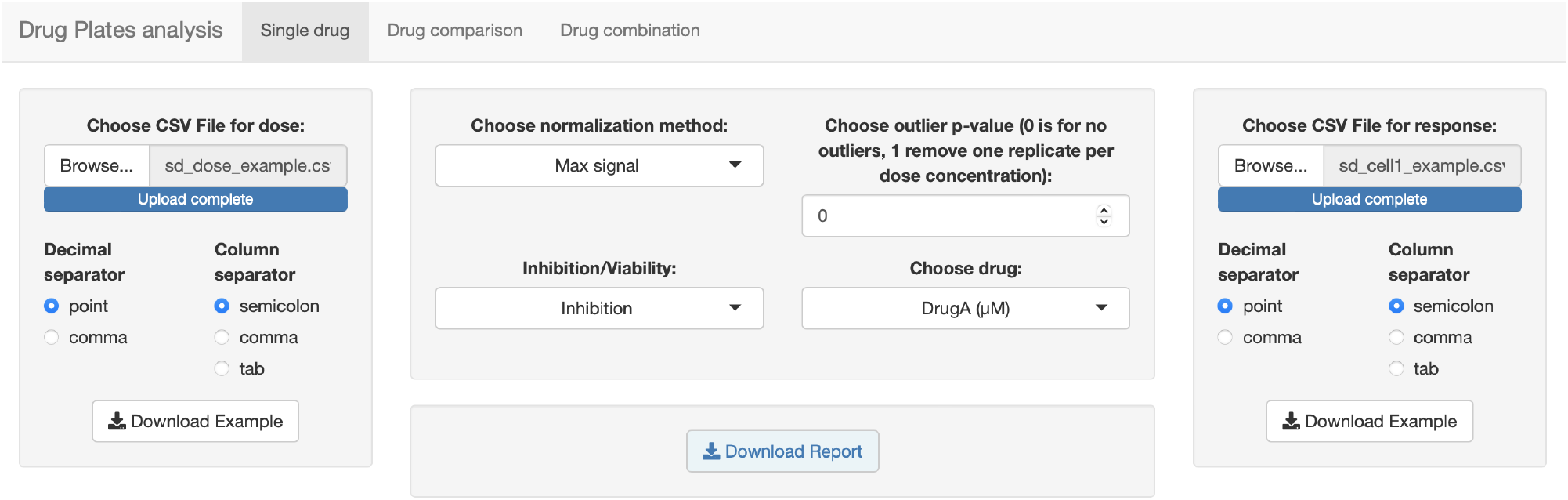
Screenshot of the first tab, dedicated to single drug analysis.

In the second tab (Figure 4) it is possible to make a comparison between different samples, such as different cell lines on which the same drugs are administered at the same doses. The input files are identical to those of the previous tab, with the difference that up to four files with drug response data can be loaded. It is thus possible to choose between the different models, also based on what has been seen in the previous tab and shows the curve corresponding to each drug. It can be represented by one to four curves, based on the number of samples you want to analyze and compare.

**Figure 4.**
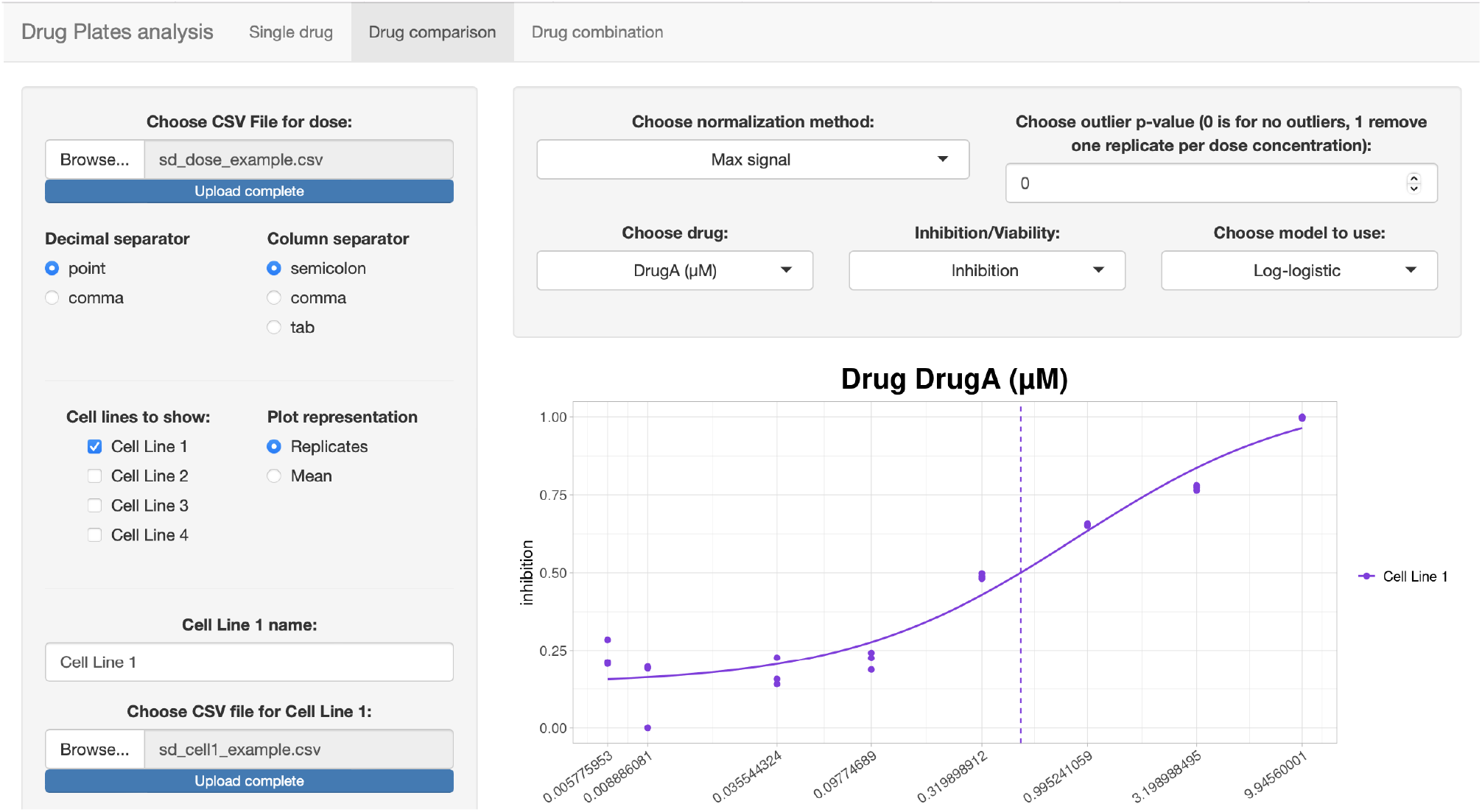
Screenshot of the second tab, dedicated to drug comparison.

Finally, in the third tab (Figure 5) we have the analysis relating to the combination of drugs. Here we have both the visualization relating to the single curves for the five models of the dose-response curve with the calculation of the R^2^, and the plots for the combination, with the representation of the CI. It is possible to choose both between the five models of the index combination and between the five of the dose-response curves, in the case of dose-effect based methods.

**Figure 5.**
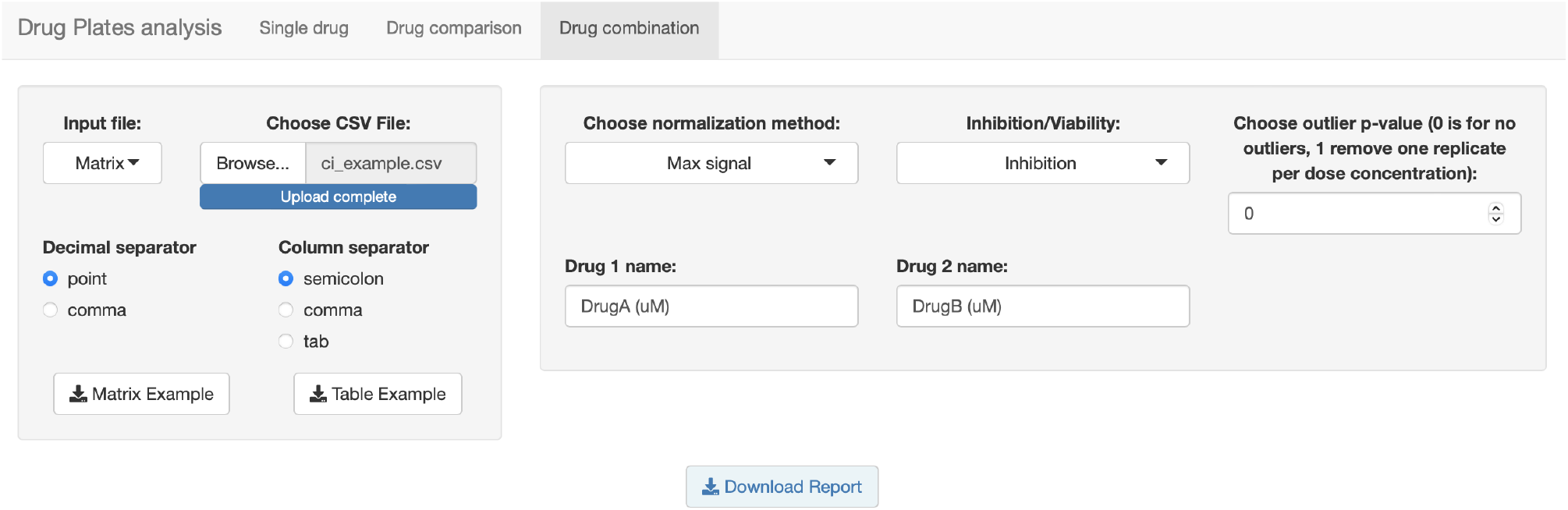
Screenshot of the third tab, dedicated to drug combination.

## Conclusions

SiCoDEA is an open-source app among the most complete (Table 2). It is developed through a Shiny interactive and easy-to-use interface (R based, like our RNA-Seq app, ARPIR [35, 36]) and it allows users to:

- Obtain the dose-response curve for different drugs by uploading a single file. For each drug, curves are displayed for all five models with relative R^2^ and IC50.
- Compare the effect of the same drug on different samples, up to a maximum of four, by choosing the most fitting model from the five options.
- Perform a combination analysis in two steps: first, visualizing the dose-response curves for the five models in the two drugs considered, second, based on the R^2^, choosing the best model to be adopted for the dose-response curve and for the CI. Results are displayed in an isobologram plot and in a heatmap.

**Table 2.** SiCoDEA offers the greatest number of options compared to the various existing tools.

|  | CI models | Drug-response models | Open source | Report | Single drug analysis | Customable outlier analysis | Platform |
| --- | --- | --- | --- | --- | --- | --- | --- |
| **SiCoDEA** | 5 | 5 | ✓ | ✓ | ✓ | ✓ | Win/Mac/Linux |
| **SynergyFinder Plus** | 4 | 1 | ✓ | ✓ | ✓ |  | Win/Mac/Linux |
| **CompuSyn** | 1 | 1 |  |  | ✓ |  | Win |
| **DDCV** | 1 | 1 | ✓ | ✓ |  |  | Win/Mac/Linux |

Data input can be easily integrated into Laboratoty Information Management Systems (LIMS) which support analysis plate readings, such as adLIMS [37]. SiCoDEA can be a useful tool, next to the already existing ones, for drug combinatorial treatments analysis as it introduces different mathematical models allowing accurate fitting of data. It is a flexible app that allows you to better adapt the analysis parameters based on the data.

## Availability and requirements

**Project name:** SiCoDEA

**Project home page:** https://github.com/giuliospinozzi/SiCoDEA

**Operating systems:** Unix (Linux, Mac)

**Programming language:** R

**Other requirements:** R >4.0, shiny, shinyjs, plyr, car, drc, ggplot2, tidyr, gplots, outliers, scales, rlist, dplyr (for standalone version available in github repository). Only a web browser for the website version.

**License:** GNU GPL

**Any restrictions to use by non-academics:** no restrictions

## Availability of data and materials

The datasets supporting the conclusions of this article are included within the article (and its additional files). The software is available in the GitHub repository: https://github.com/giuliospinozzi/SiCoDEA.

## Authors’ contributions

GS conceived of the study, participated in its design and coordination. GS and VT created the application (SiCoDEA) and wrote the manuscript. AF, IG, RR generated the experiments with AML cell lines. FM, SP, SD and SS supported the work. GS, MPM and BF coordinated all work. All authors read and approved the final manuscript.

## Ethics approval and consent to participate

Not applicable.

## Consent for publication

Not applicable.

## List of additional data files

Additional file 1: **Supplementary Material.** Read me and user guide for SiCoDEA.

## References

1. Frei E, 3rd, Karon M, Levin RH, Freireich EJ, Taylor RJ, Hananian J, Selawry O, Holland JF, Hoogstraten B, Wolman IJ et al: The effectiveness of combinations of antileukemic agents in inducing and maintaining remission in children with acute leukemia. Blood 1965, 26(5):642–656.

2. DeVita VT, Jr., Chu E: A history of cancer chemotherapy. Cancer Res 2008, 68(21):8643–8653.

3. Zhou F, Zhao W, Gong X, Ren S, Su C, Jiang T, Zhou C: Immune-checkpoint inhibitors plus chemotherapy versus chemotherapy as first-line treatment for patients with extensive-stage small cell lung cancer. J Immunother Cancer 2020, 8(2).

4. Mullard A: 2021 FDA approvals. Nat Rev Drug Discov 2022, 21(2):83–88.

5. [https://www.nytimes.com/2017/08/12/health/cancer-drug-trials-encounter-a-problem-too-fewpatients.html]

6. Hay M, Thomas DW, Craighead JL, Economides C, Rosenthal J: Clinical development success rates for investigational drugs. Nat Biotechnol 2014, 32(1):40–51.

7. Moore TJ, Zhang H, Anderson G, Alexander GC: Estimated Costs of Pivotal Trials for Novel Therapeutic Agents Approved by the US Food and Drug Administration, 2015-2016. JAMA Intern Med 2018, 178(11):1451–1457.

8. Foucquier J, Guedj M: Analysis of drug combinations: current methodological landscape. Pharmacol Res Perspect 2015, 3(3):e00149.

9. Slinker BK: The statistics of synergism. J Mol Cell Cardiol 1998, 30(4):723–731.

10. Lehar J, Zimmermann GR, Krueger AS, Molnar RA, Ledell JT, Heilbut AM, Short GF, 3rd, Giusti LC, Nolan GP, Magid OA et al: Chemical combination effects predict connectivity in biological systems. Molecular systems biology 2007, 3:80.

11. Bliss CI: The toxicity of poisons applied jointly. Annals of Applied Biology 1939.

12. Loewe SM, H. : Über Kombinationswirkungen. Naunyn-Schmiedebergs Archiv für experimentelle Pathologie und Pharmakologie volume 1926.

13. Loewe S: Die quantitativen Probleme der Pharmakologie. Ergebnisse Der Physiologie 1928.

14. Loewe S: The problem of synergism and antagonism of combined drugs. Arzneimittelforschung 1953, 3(6):285–290.

15. Yadav B, Wennerberg K, Aittokallio T, Tang J: Searching for Drug Synergy in Complex Dose-Response Landscapes Using an Interaction Potency Model. Comput Struct Biotechnol J 2015, 13:504–513.

16. Yadav B, Wennerberg K, Aittokallio T, Tang J: Corrigendum to “Searching for drug synergy in complex dose-response landscapes using an interaction potency model” [Comput. Struct. Biotechnol. J. 13 (2015) 504-513]. Comput Struct Biotechnol J 2017, 15:387.

17. Chou TC, Talalay P: Quantitative analysis of dose-effect relationships: the combined effects of multiple drugs or enzyme inhibitors. Adv Enzyme Regul 1984, 22:27–55.

18. Chou TC: Drug combination studies and their synergy quantification using the Chou-Talalay method. Cancer Res 2010, 70(2):440–446.

19. Zheng S, Wang W, Aldahdooh J, Malyutina A, Shadbahr T, Pessia A, Jing T: SynergyFinder Plus: towards a better interpretation and annotation of drug combination screening datasets. bioRxiv 2021.

20. Zhang T: Drug-Drug Combination Visualization (DDCV): Evaluation of Drug-Drug Interactions using Shiny by RStudio. 2015.

21. Martelli MP, Gionfriddo I, Mezzasoma F, Milano F, Pierangeli S, Mulas F, Pacini R, Tabarrini A, Pettirossi V, Rossi R et al: Arsenic trioxide and all-trans retinoic acid target NPM1 mutant oncoprotein levels and induce apoptosis in NPM1-mutated AML cells. Blood 2015, 125(22):3455–3465.

22. shiny: Web Application Framework for R. R package version 1.1.0. [https://cran.r-project.org/web/packages/shiny/index.html]

23. shinyjs: Easily Improve the User Experience of Your Shiny Apps in Seconds [https://cran.r-project.org/web/packages/shinyjs/index.html]

24. Wickham H: The Split-Apply-Combine Strategy for Data Analysis. Journal of Statistical Software 2011.

25. An R Companion to Applied Regression [https://socialsciences.mcmaster.ca/jfox/Books/Companion/]

26. Ritz C, Baty, F., Streibig, J. C., Gerhard, D. : Dose-Response Analysis Using R. PLoS ONE 2015.

27. ggplot2: Elegant Graphics for Data Analysis

28. tidyr: Tidy Messy Data [https://CRAN.R-project.org/package=tidyr]

29. Gregory R. Warnes BB, Lodewijk Bonebakker, Robert Gentleman, Wolfgang Huber, Andy Liaw, Thomas Lumley, Martin Maechler, Arni Magnusson, Steffen Moeller, Marc Schwartz and Bill Venables gplots: Various R Programming Tools for Plotting Data. 2020.

30. Komsta L: outliers: Tests for outliers. 2011.

31. scales: Scale Functions for Visualization [https://CRAN.R-project.org/package=scales]

32. rlist: A Toolbox for Non-Tabular Data Manipulation [https://CRAN.R-project.org/package=rlist]

33. dplyr: A Grammar of Data Manipulation [https://CRAN.R-project.org/package=dplyr]

34. Grubbs FE: Sample Criteria for testing outlying observations. Ann Math Stat 1950, 21:27–58.

35. Spinozzi G, Tini V, Adorni A, Falini B, Martelli MP: ARPIR: automatic RNA-Seq pipelines with interactive report. BMC bioinformatics 2020, 21(Suppl 19):574.

36. Spinozzi G, Tini V, Mincarelli L, Falini B, Martelli MP: A comprehensive RNA-Seq pipeline includes meta-analysis, interactivity and automatic reporting. PeerJ Preprints 2018.

37. Calabria A, Spinozzi G, Benedicenti F, Tenderini E, Montini E: adLIMS: a customized open source software that allows bridging clinical and basic molecular research studies. BMC bioinformatics 2015, 16 Suppl 9:S5.

