## Additional File 1 for "SiCoDEA: a simple, fast and complete app for analyzing the effect of individual drugs and their combinations"

SiCoDEA (Single and Combined Drug Effect Analysis) is a simple, fast and complete app for analyzing the effect of individual drugs and their combinations and is available at: https://sicodea.shinyapps.io/shiny/. The source code is available in our GitHub repository: https://github.com/giuliospinozzi/SiCoDEA. A video tutorial is also available: https://youtu.be/kzzdU83r0_I. The app consists of three panels, each of which allows you to carry out a different analysis: a single drug analysis, a comparison of different drugs or an analysis of drug combinations.

A step-by-step guide of all the analyzes that can be carried out with SiCoDEA is provided below.

#### Single drug analysis

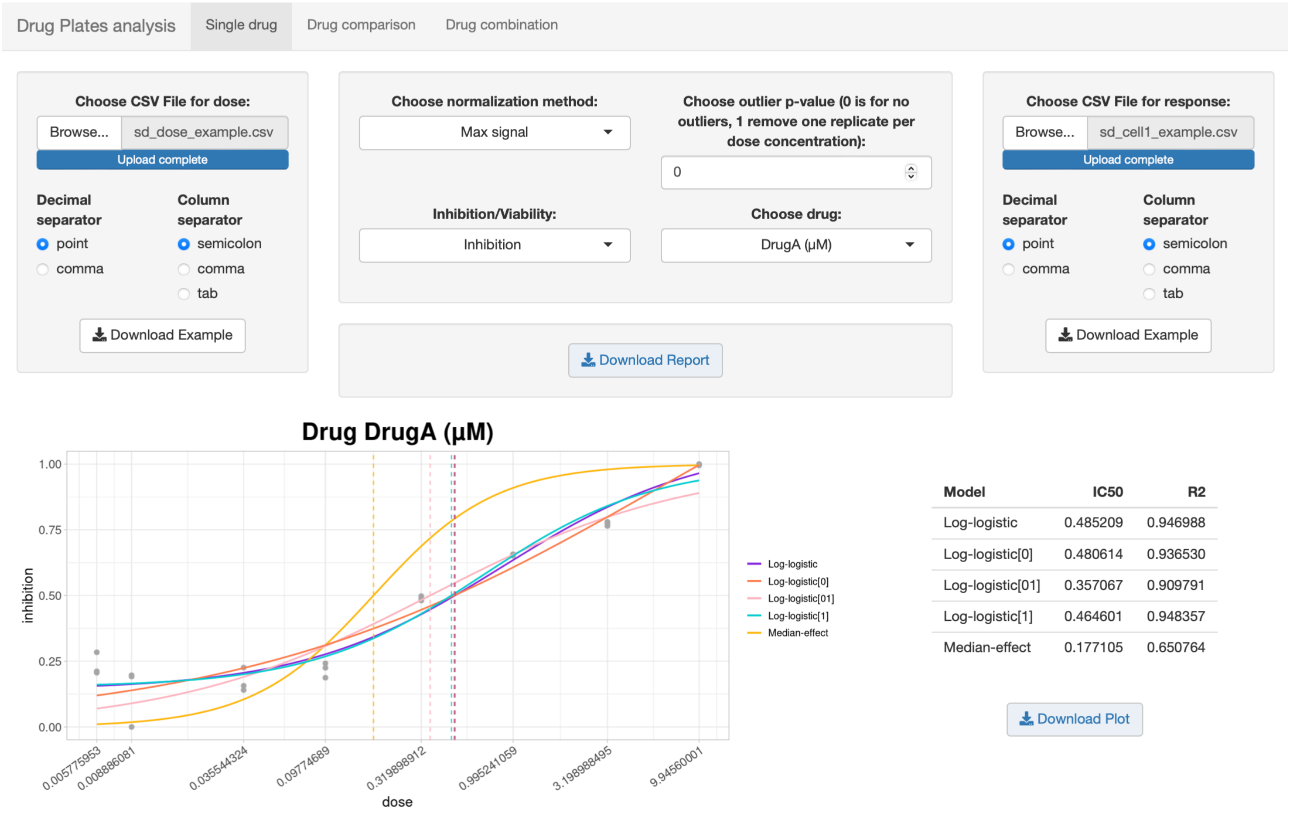


It can be done in the first tab, called "Single drug".

1. Upload files. The data relating to drug doses and responses must be uploaded to the two side panels. To get an idea of the required formatting, you can download the sample files. For the dose file you need a matrix in which the first column contains the names of the different drugs under examination and in which each row contains the sequence of doses used for the relative drug. For the second file, however, a matrix with the response values calculated for the different doses is required. The sequence and position in the response file must be

corresponding to the dose values represented in the other file. Both files must be in CSV format, but you can choose whether to use comma, semicolon or tab separators.

2. Choose options. Once the files have been uploaded, it is then possible to choose different options in the central panel.
   a. Normalization can be performed considering the maximum signal of the entire plate as a basis, or considering the average of the replicates obtained for drug concentrations equal to zero as the baseline. If, on the other hand, the normalization has already been carried out on the loaded data, you can choose the "no normalization" option.
   b. Based on the experiment in question, it is then possible to choose between inhibition and viability to indicate the information provided by the response values loaded.
   c. A Grubbs' test is performed on the replicates to assess the presence of any outliers and the filter applied is based on the calculated p-value. It can be set in the central panel using values ranging from 0 to 1, where in the first case replicas are not eliminated while in the second one is eliminated for each dose concentration. Rather than setting a fixed value, we preferred to leave the possibility of choice in order to evaluate in real time and visually from the plot the actual nature of an outlier.
   d. Finally, there is a drop-down menu that allows you to navigate between the different drugs present in the input file.

3. Plot generation. The plot below shows, for each drug, the individual values at the different doses as points, while the curves represent the expected trend of the five different models examined. For each, the value of the IC50 is also shown with a dashed vertical line. The next table summarizes the value of each IC50, as well as the value of $R^2$ for each model, so as to be able to evaluate the most suitable for the data being analyzed. The closer to 1 the value of $R^2$, the better the model for the data under consideration.

4. Download plot. Finally, there is a "Download plot" button that allows you to download the single plot that is displayed at that moment, or the "Download report" button, which generates a report containing all the plots for the various drugs.

#### Comparison of different drugs

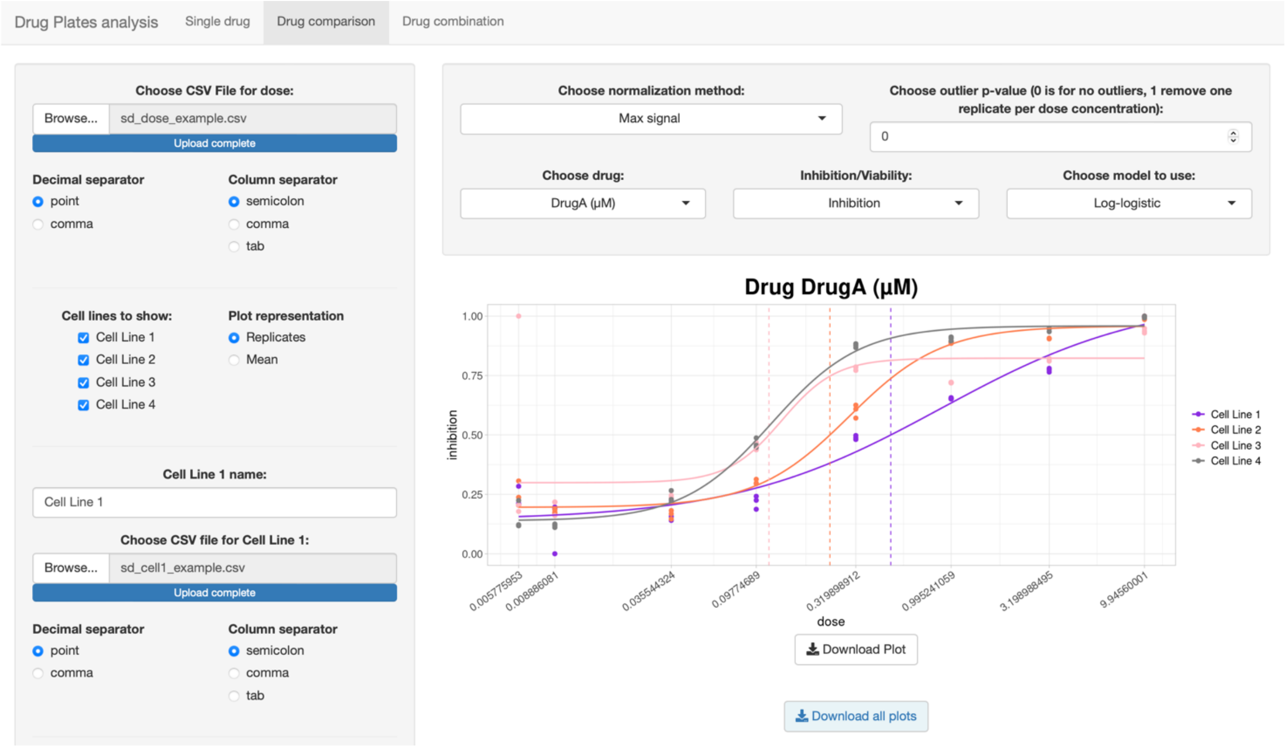


The second tab concerns the comparison between drugs.

1. Upload files. Again, there is a side panel for uploading files. The formatting required for doses and responses is identical to that of the previous tab. In this case, however, it is possible to load up to four files for the response values, while the dose file remains the same; for example, these may be four different cell lines to which the same sequence of doses of a drug is applied to evaluate its effectiveness.

   It is possible to choose which and how many cell lines to load and show by selecting the different ticks and with each new tick selected a new panel will appear to load the file. You can also choose whether to represent replicates as distinct points in the plot or as mean and standard deviation.

   In this case also, as in the previous one, the names of the drugs are those reported in the input file, while the names of the cell lines can be assigned in the specific spaces.

2. Choose options. The options in the central panel are the same as those seen in the previous tab. However, we also find the option relating to the model to be used.

3. Plot generation. In this case, in fact, in the plot the different cell lines are compared and not the different models and therefore a specific model must be chosen to be represented also, possibly, on the basis of the $R^2$ calculated in the previous tab.

4. Download plot.

#### Analysis of drug combinations

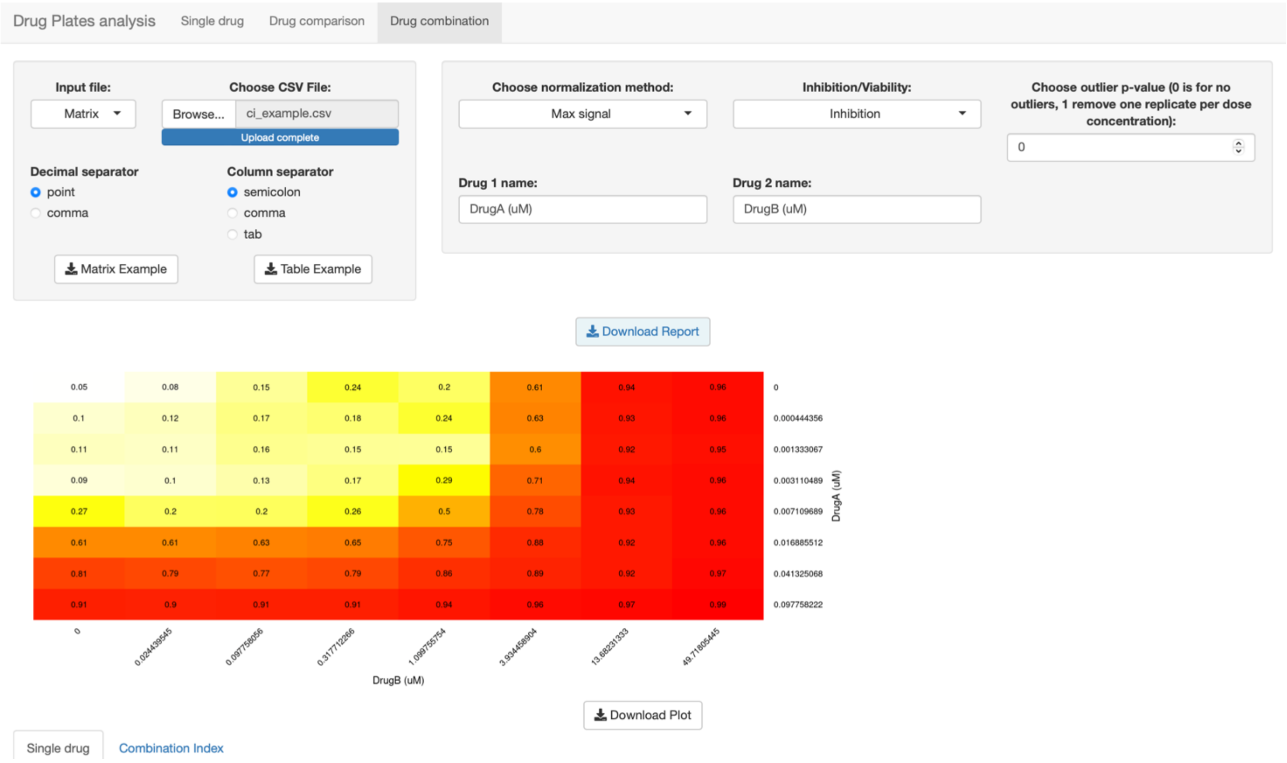


Finally, in the third tab we have the analysis of drug combinations.

1. Upload files. It is possible to choose in which format to load the input file, whether as a table, showing a column with the doses of a drug, a column with the doses of the other drug and a column with the recorded values, or as a matrix, reporting the doses of a drug in the first column, those of the other drug in the first row and the values recorded in the central cells. There must also be doses of one drug and the other equal to zero in order to calculate the baseline value and build the curve of the individual drugs. For clarity, there is a downloadable example for each format.

2. Choose options. The options for the normalization method, for the choice between viability and inhibition, for the p-value of the outliers are the same as already seen. In addition, it is also possible to change the name of the drugs directly from the interface.

3. Plot generation. The first plot shows a matrix with the average of the replicates present in the source file for each dose combination and there is the usual button to download it. Below then we have two tabs, one for the plots relating to the individual drugs and the other for the combination.

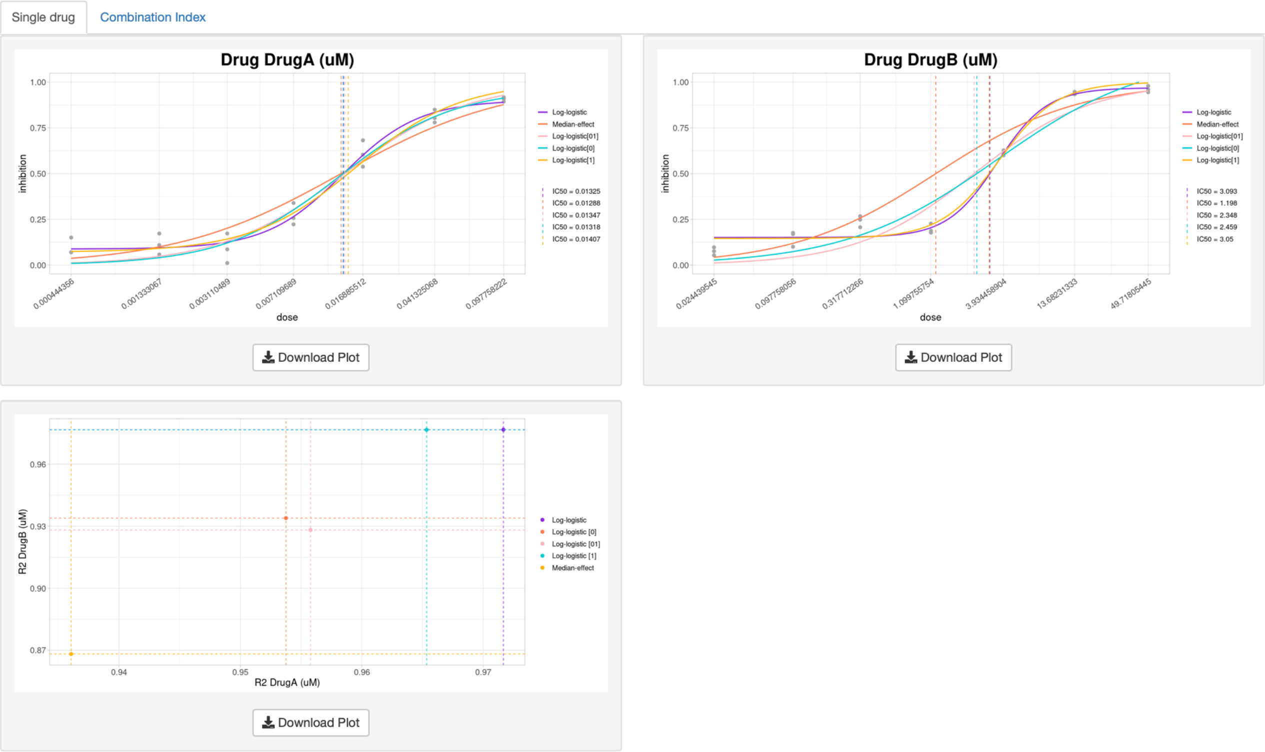


a. Single drug plots. In the first tab we have the plots for the dose-response curve of the two drugs under examination and the representation of all five models. This is the same plot also seen in the "Single Drug" tab. In this case also, the $R^2$ value is calculated for each of the five models and in both drugs. The results are shown in the last plot, in which there are the points of intersection of the $R^2$ for one drug and the other. The points that are most located at the top right are those with the most suitable model for the data under examination. This way we can choose the right model for our experiment.

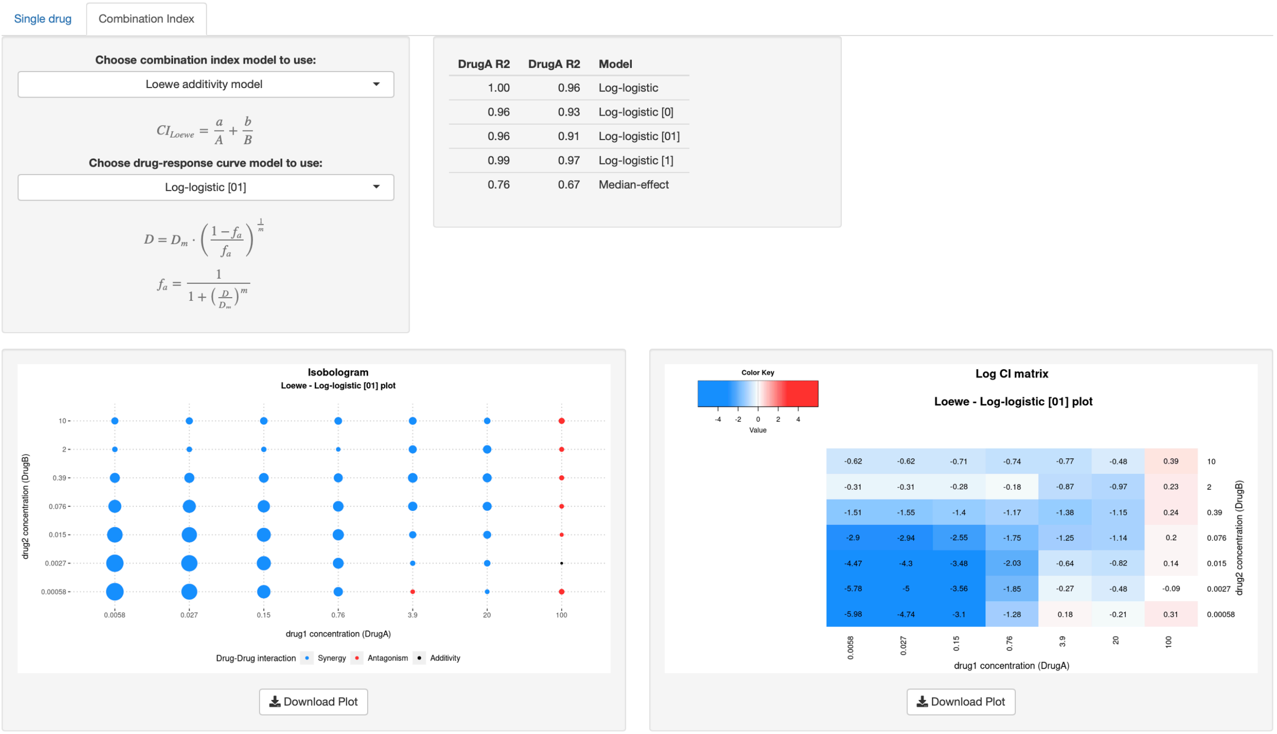


b. Combination index plots. In the second tab it is possible to choose the model for the combination index from five different options: *Response Additivity* model [1], *Highest Single Agent* (HSA) model [2], *Bliss Independence* model [3], *Loewe Additivity* model [4-6] and *Zero Interaction Potency* (ZIP) model [7, 8]. By selecting the different models from the drop-down menu, the corresponding formula will appear in the panel. In the case of the Loewe Additivity model and ZIP model, since they use an Effect-Based Strategy, it is also necessary to choose the model for the dose-response curve. The next table shows the two $R^2$ values for the five models, the same ones also present in the previous plot. The two plots below show the values of the combination index for each combination of doses under examination based on the models chosen; blue represents synergy, red antagonism and black additivity. These are two different representations of the same results: in the first case there are dots with dimensions proportional to the strength of synergy or antagonism, while in the second case we have a matrix with colors of intensity proportional to the values of combination index reported within the cells. You can then browse through the various models to see how the results vary for the same data.

4. Download plots. Finally, you can download the single plots or the report with all the plots.

### Reference


1. Slinker BK: **The statistics of synergism**. *J Mol Cell Cardiol* 1998, **30**(4):723-731.
2. Lehar J, Zimmermann GR, Krueger AS, Molnar RA, Ledell JT, Heilbut AM, Short GF, 3rd, Giusti LC, Nolan GP, Magid OA *et al*: **Chemical combination effects predict connectivity in biological systems**. *Molecular systems biology* 2007, **3**:80.
3. Bliss CI: **The toxicity of poisons applied jointly**. *Annals of Applied Biology* 1939.
4. Loewe SM, H. : **Über Kombinationswirkungen**. *Naunyn-Schmiedebergs Archiv für experimentelle Pathologie und Pharmakologie volume* 1926.
5. Loewe S: **Die quantitativen Probleme der Pharmakologie**. *Ergebnisse Der Physiologie* 1928.
6. Loewe S: **The problem of synergism and antagonism of combined drugs**. *Arzneimittelforschung* 1953, **3**(6):285-290.
7. Yadav B, Wennerberg K, Aittokallio T, Tang J: **Searching for Drug Synergy in Complex Dose-Response Landscapes Using an Interaction Potency Model**. *Comput Struct Biotechnol J* 2015, **13**:504-513.
8. Yadav B, Wennerberg K, Aittokallio T, Tang J: **Corrigendum to "Searching for drug synergy in complex dose-response landscapes using an interaction potency model" [Comput. Struct. Biotechnol. J. 13 (2015) 504-513]**. *Comput Struct Biotechnol J* 2017, **15**:387.
